# Rescue of Auditory Function by a Single Administration of AAV-*TMPRSS3* Gene Therapy in Aged Mice of Human Recessive Deafness DFNB8

**DOI:** 10.1101/2023.02.25.530035

**Authors:** Wan Du, Volkan Ergin, Corena Loeb, Mingqian Huang, Stewart Silver, Ariel Miura Armstrong, Zaohua Huang, Channabasavaiah B. Gurumurthy, Hinrich Staecker, Xuezhong Liu, Zheng-Yi Chen

## Abstract

Patients with mutations in the *TMPRSS3* gene suffer from recessive deafness DFNB8/DFNB10 for whom cochlear implantation is the only treatment option. Poor cochlear implantation outcomes are seen in some patients. To develop biological treatment for TMPRSS3 patients, we generated a knock-in mouse model with a frequent human DFNB8 *TMPRSS3* mutation. The *Tmprss3*^*A306T/A306T*^ homozygous mice display delayed onset progressive hearing loss similar to human DFNB8 patients. Using AAV2 as a vector to carry a human *TMPRSS3* gene, AAV2-h*TMPRSS3* injection in the adult knock-in mouse inner ears results in *TMPRSS3* expression in the hair cells and the spiral ganglion neurons. A single AAV2-h*TMPRSS3* injection in aged *Tmprss3*^*A306T/A306T*^ mice leads to sustained rescue of the auditory function, to a level similar to the wildtype mice. AAV2-h*TMPRSS3* delivery rescues the hair cells and the spiral ganglions. This is the first study to demonstrate successful gene therapy in an aged mouse model of human genetic deafness. This study lays the foundation to develop AAV2-h*TMPRSS3* gene therapy to treat DFNB8 patients, as a standalone therapy or in combination with cochlear implantation.

## Introduction

Hearing loss (HL) is one of the most common sensory deficit disorders that affect about 466 million people.^1^ Expected to afflict one in ten individuals by 2050, HL poses a social and emotional toll, and a growing worldwide annual economic toll.^1^ HL has been linked to increased instances of social isolation and higher risk for dementia and depression.^2,3^ Genetic hearing loss affects one in 500 newborns. There are no treatments available to reverse or prevent genetic deafness.^4,5^ Currently hearing aids and cochlear implants are the only treatment options that require residual hearing function or the ability to stimulate the cochlear nerve, respectively. While genetic hearing loss patients can benefit from cochlear implantation (CI), those patients with ganglion deficits will be left without any treatment option.

Rapid progress has been made in the understanding of the genetic etiology of human HL.^6^ Genetic testing and diagnosis of HL provide essential information for further genetic therapies.^7,8^ Adeno-Associated Virus (AAV), a non-replicative viral vector with low immunogenicity and little ototoxicity, is one of the most promising gene therapy tools for transducing broad cellular tropism.^9,10^ Gene therapies including gene editing which can replace, edit, or silence genes offer a promising avenue for treatment. Among them, gene replacement is appropriate for treating recessive monogenic disorders and has achieved the most clinical success, such as Luxturna^11^ for an inherited retinal disease Leber’s congenital amourosis (LCA) and Zolgensma^12^ for spinal muscular atrophy (SMA). As the inner ear is an isolated organ that can be accessed safely by local injection, a number of gene replacement and over-expression studies targeting hearing loss have been conducted successfully, resulting in hearing rescue and the survival of hair cells or spiral ganglion neurons.^13-35^ However, with the exception of *Otof*^25^, all the studies were performed in neonatal animals, which raises the question of the suitability of the approach in fully mature adult inner ear.

TMPRSS3 protein, a type II transmembrane serine protease, is necessary for normal hearing in mammals. *TMPRSS3* gene mutations account for approximately 12-13% of HL families which are negative for common genetic mutations.^36,37^ Two different phenotypes were present in individuals with *TMPRSS3* mutations: prelingual (DFNB10, OMIM 605511) and the delayed onset and postlingual (DFNB8, OMIM 601072).^38^ Previous study has shown TMPRSS3 as a necessary permissive factor for cochlear hair cell activation and survival upon the onset of hearing.^39^ Mice with truncated *TMPRSS3* protease domains of mutation *TMPRSS3*^Y260X^ showed congenital hearing loss, and rapidly loss of hair cells and progressive loss of spiral ganglion neurons.^39^

To develop a mouse model with a TMPRSS3 mutation suitable for gene therapy intervention, we constructed a DFNB8 mouse model with a knock-in (KI) of a human *TMPRSS3* mutation (c.916G>A, Ala306⟶Thr), which causes adulthood onset and progressive recessive HL. We show by injection in older mice, a human *TMPRSS3* gene carried by an AAV2 is re-expressed in the hair cells and the modiolus region. *TMPRSS3* inner ear delivery rescues hearing in aged KI mice, concomitant with the survival of hair cells and spiral ganglion neurons.

## Results

### Generation of *Tmprss3* p.A306T knock-in mouse model

*TMPRSS3* gene has 13 exons and encodes a protein which consists of 453 amino acids, containing four domains (**Figure 1A**). A human *TMPRSS3* mutation in Ala306 identified in hearing loss patients is associated with DFNB8/10.^40-44^ Ala306 is highly conserved across species from zebrafish to human (**Figure 1B**). We chose to use the inbred strain of CBA/CaJ to create a KI mouse model as CBA/CaJ does not suffer from age-related hearing loss as in the strain of C57BL/6J and will allow us to analyze the rescue effect over time.^45^ We used CRISPR/Cas9 technology^46-49^ to create a KI mutation c.916G>A, which changed Ala to Thr at the amino acid position 306 (p.A306T). In brief, to introduce mutations of Tmprss3 c.916G>A; 918 C>T, i.e. p.Ala306Thr, ribonucleoprotein (RNP) complex of Cas9 protein, UGGCUCGGACAGCUUCAUGA guide RNA with the donor oligonucleotide of ACAGCAAGTACAAGCCAAAGCGGCTGGGCAATGACATAACTC TCATGAAGCTTTCCGAGCCACTCACCTTTGACGAGACCATCCAGCC were injected into CBA/CaJ zygotes (Jackson laboratory stock number 000656). The microinjected zygotes were then transferred into pseudo-pregnant females.^49^ The target genomic DNA region from the founder were amplified and sequenced using PCR primer pairs of Tmprss3F, GGAGATCCCACATCTCTCACC; and Tmprss3R, AAATGCTATGCACCTACATCAAC. After the mutations were confirmed by sequencing, founder mouse carrying the mutations were mated to generate wildtype (WT), heterozygous (Het) and homozygous (Homo) mutation mice. A representative genotyping mice determined by Sanger sequencing is shown (**Figure 1C**).

**Figure. 1.**
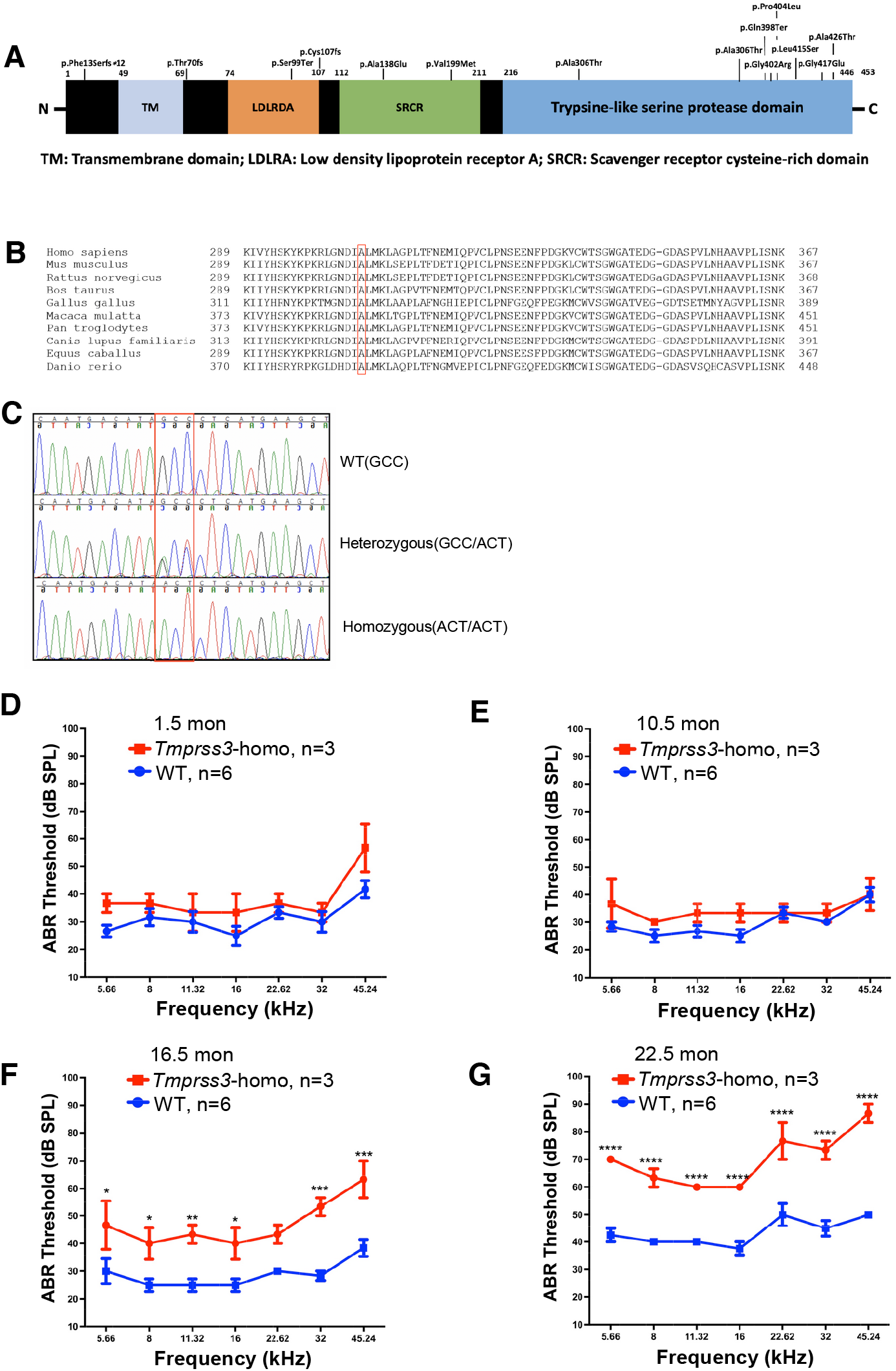
Generation of *Tmprss3* p.A306T knock-in mouse model with late-onset progressive hearing loss. (A) Summary of human the TMPRSS3 protein domains and mutations. (B) Conservation of the 306 alanine from in species human to zebrafish. (C) Sanger sequencing of *TMPRSS3* c.916G>A; c.918C>T mutant mice. (D-G) ABR thresholds in *Tmprss3*^*A306T/A306T*^ homozygous mice ears (red), compared to wild-type ears (blue) at 1.5 months (D), 10.5 months (E), 16.5 months (F), and 22.5 months (G), respectively. Significant hearing loss by the elevated ABR thresholds was seen at 16.5 months, which became more severe at 22.5 months. Values and error bars reflect mean ± SEM. Statistical analyses were performed by two-way ANOVA with Bonferroni correction for multiple comparisons: P value style, <0.05 (*), <0.01 (**), <0.001(***), <0.0001(****).

### *Tmprss3*^A306T/A306T^ mice display late-onset progressive hearing loss

With the aim of rescuing auditory function in the *Tmprss3*^*A306T/A306T*^ mice, we first performed auditory brain stem responses (ABR, represent sound-evoked neural output of the cochlea) and distortion product otoacoustic emissions (DPOAE, measurement of outer hair cell function) tests to study hearing in *Tmprss3*^*A306T/A306T*^ mice (**Figure 1D-G and S1**). We found that there was no change in ABR/DPOAE thresholds or ABR wave1 amplitudes in the *Tmprss3*^*A306T/A306T*^ ears compared to the WT CBA/CaJ ears across all frequencies at 1.5 months and 10.5 months (**Figure 1D-E and S1A-D**). By 16.5 months, the ABR thresholds were significantly elevated in the *Tmprss3*^*A306T/A306T*^ ears by an average of 15 dB compared to WT ears across most frequencies (**Figure 1F**). DPOAE thresholds were significantly elevated at 11 and 45 kHz and wave 1 amplitudes showed significant reductions at 32 kHz with 80- and 90-dB SPL stimulation (**Figure S1E, F**). At 22.5 months, ABR thresholds were significantly elevated by an average of 26 dB across all frequencies (**Figure 1G**). We did not find changes in wave1 amplitudes in *Tmprss3*^*A306T/A306T*^ ears compared to WT ears at 22.5 months (**Figure S1H**), as both were much lower than that at 16.5 months (**Figure S1F**), an indication that neuronal activates were reduced in the WT mice due to aging. The DPOAE thresholds were elevated in *Tmprss3*^*A306T/A306T*^ ears compared to WT ears at 22.5 months (**Figure S1G**). These results showed late-onset progressive hearing loss that started after 10.5 months with significantly elevated ABR/DPOAE thresholds and reduction in wave 1 amplitudes at 16.5 months, which further deteriorated by 22.5 months of age.

### Designing an AAV-*TMPRSS3* gene therapy strategy

The phenotype of late onset and progressive hearing loss in the *Tmprss3*^*A306T/A306T*^ KI mice supports the development of a gene therapy strategy to rescue hearing in adult mouse cochlea which is relevant to humans clinically. We chose to use AAV2 for delivery to evaluate gene therapy in the *Tmprss3*^*A306T/A306T*^ mice as AAV2 transduces the inner hair cells (IHCs) and the outer hair cells (OHCs) with high efficiency: AAV2 transduced all the IHCs and a majority of OHCs in the apex and mid-turn with a slight reduction in the base turn in adult mouse cochlea (**Figure S2**). The size of *TMPRSS3* gene is less than 1.5 kb, making it ideal for AAV mediated delivery strategy. We cloned the coding sequences of mouse-*Tmprss3* into the AAV2 backbone to produce AAV2-*Tmprss3*. To evaluate potential functional ototoxicity of AAV2-*Tmprss3*, we injected the AAV2-*Tmprss3* in adult WT CBA/CaJ and tested hearing by ABR and DPOAE four weeks later. To our surprise, we found that ABR thresholds were significantly elevated by an average of 35 dB across all frequencies in AAV2-*Tmprss3* injected ears compared to uninjected ears (**Figure 2A**). Similarly, there was a complete loss of DPOAE across frequencies in the injected ears (**Figure 2B**). Significant ABR threshold shifts and the loss of DPOAE indicated a major damage by AAV2-*Tmprss3* to the mouse inner ear. To study the cellular ototoxicity by AAV2-*Tmprss3*, we quantified the number of OHC and IHC by whole-mount labeling of AAV2-*Tmprss3* injected ears and uninjected contralateral ears. There was a slight reduction in IHCs in AAV2-*Tmprss3* injected ears compared to uninjected ears, especially in the middle and basal turns (**Figure 2C, E**). There was a major reduction in the number of OHCs across the entire cochlear turn with the base turn being most severely affected in AAV2-*Tmprss3* injected ears (**Figure 2D, F**). These results demonstrated that AAV2-*Tmprss3* induced ototoxicity by damaging especially OHCs, resulting in profound hearing loss.

**Figure. 2.**
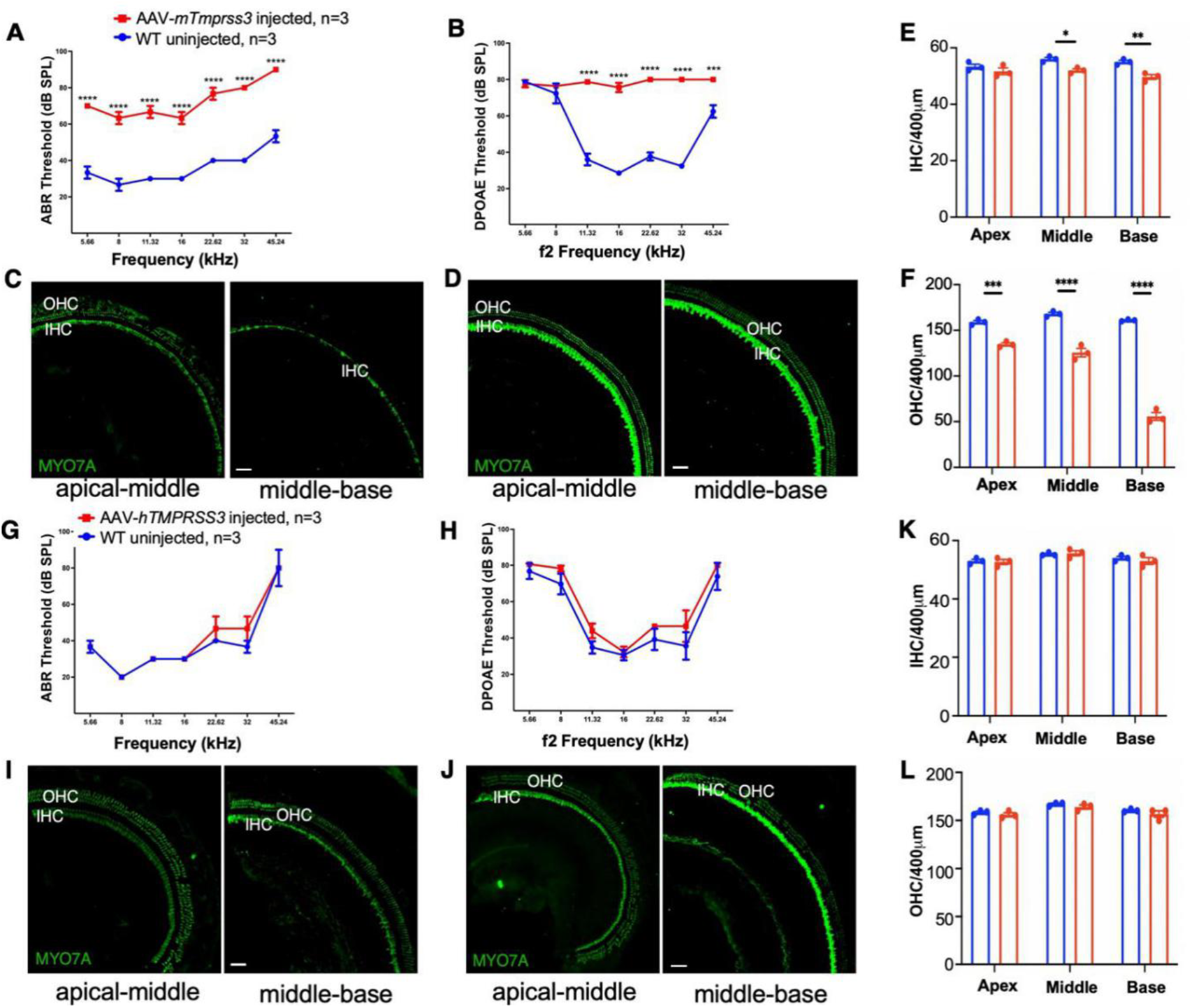
AAV2-h*TMPRSS3* gene therapy design strategy. (A) ABR thresholds were significantly elevated in WT mice ears injected with AAV2-*Tmprss3* (red), compared to uninjected ears (blue) at 2 months. (B) DPOAE thresholds were no longer detectable in WT mice ears injected with AAV2-*Tmprss3* (red), compared to uninjected ears (blue) at 2 months. (C) In an injected WT cochlea, major OHC loss and some IHC loss were seen in the middle-base turn, whereas some OHC loss was seen in the apical-middle turn. MYO7A labels hair cells. (D) In the uninjected contralateral ear, a full set of OHC and IHC were seen along the cochlear turns. Scale bar, 50μm. (E-F) Quantification of IHCs (E) and OHCs (F) per 400-μm section for three injected (red) and uninjected (blue) cochleae from three mice at 2 months. (G) ABR thresholds in WT mice ears injected with AAV2-h*TMPRSS3* (red), compared to uninjected ears (blue) at 3 months. (H) DPOAE thresholds in WT mice ears injected with AAV2-h*TMPRSS3* (red), compared to uninjected ears (blue) at 3 months. (I) A full set of OHC and IHCs were seen in an AAV2-h*TMPRSS3* injected WT ear along the cochlear turn. (J) The contralateral uninjected inner ear showed a full set of OHC and IHC. Scale bar, 50μm. (K-L) Quantification of IHCs (K) and OHCs (L) per 400-μm section for three injected (red) and uninjected (blue) cochleae from three mice at 3 months. Values and error bars reflect mean ± SEM. Statistical analyses were performed by two-way ANOVA with Bonferroni correction for multiple comparisons: P value style, <0.05 (*), <0.01 (**), <0.001(***), <0.0001(****).

While we did not know the origin of the mouse *Tmprss3* mediated toxicity, one approach to circumvent the issue is to test the use of the human *TMPRSS3* (h*TMPRSS3*) gene for delivery, as the human *TMPRSS3* gene will be used in the clinic. We constructed an AAV2-h*TMPRSS3* and injected into the adult WT CBA/CaJ inner ears and performed hearing study and inner ear characterization four weeks later. In the injected ears, ABR threshold showed a similar profile to uninjected inner ears (**Figure 2G**). DPOAE thresholds in the injected ears were slightly elevated but was not significant from that of uninjected ears (**Figure 2H**). There was no change in the number of IHCs and OHCs in AAV2-h*TMPRSS3* injected and uninjected ears (**Figure 2I-L**). Taken together, we conclude that the AAV2-h*TMPRSS3* inner ear delivery did not impair normal hearing or damage hair cells, and thus AAV2-h*TMPRSS3* was suitable for gene therapy in the *Tmprss3*^*A306T/A306T*^mouse model.

### AAV2-mediated *TMPRSS3* expression in HC and SGN of mouse cochlea

The expression of *Tmprss3* gene or the localization of the TMPRSS3 protein is not well studied in adult mouse inner ear. To study the expression pattern of *Tmprss3* in adult mouse inner ear and to evaluate the expression of the human *TMPRSS3* gene as the result of AAV2-mediated delivery, we performed RNA-fluorescence *in situ* hybridization (RNAscope)^50,51^ to detect mouse *Tmprss3* and human *TMPRSS3* gene, respectively. In WT adult mouse inner ear, a probe against the mouse *Tmprss3* mRNA showed strong hybridization to the endogenously *Tmprss3* mRNA that was primarily in the cochlear hair cells (MYO7A^+^) in the sensory epithelium (**Figure 3A**). We then used a human *TMPRSS3* probe for RNAscope in WT adult mouse inner ear and did not detect significant signals above the background (**Figure 3B**). We quantified the hybridization signals by ImageJ. In WT adult inner ear, the endogenous *Tmprss3* mRNA was expressed at a higher level in the IHCs than the OHCs, whereas no human *TMPRSS3* signals was detected (**Figure 3E**). Our data showed the human *TMPRSS3* probe does not cross-react to the mouse *Tmprss3* mRNA, demonstrating the specificity of the human *TMPRSS3* probe. We then injected AAV2-*hTMPRSS3* into adult *Tmprss3*^*A306T/A306T*^ mouse inner ear and performed RNAscope one month after injection. RNAscope showed *TMPRSS3* mRNA signals that became clearly visible in the hair cells (**Figure 3C**) compared to uninjected cochlea where no signal was detected (**Figure 3D**) of the same animals by the same probe. Quantification showed a similar level of human *TMPRSS3* transgene was detected in the IHC and OHC of injected *Tmprss3*^*A306T/A306T*^ inner ears (**Figure 3E**), whereas the overall *TMPRSS3* transgene expression level was lower than the endogenous *Tmprss3* mRNA in WT inner ear (**Figure 3A, C, and E**). This study confirmed species-specific probes for mRNA analyses by the RNAscope method.

**Figure. 3.**
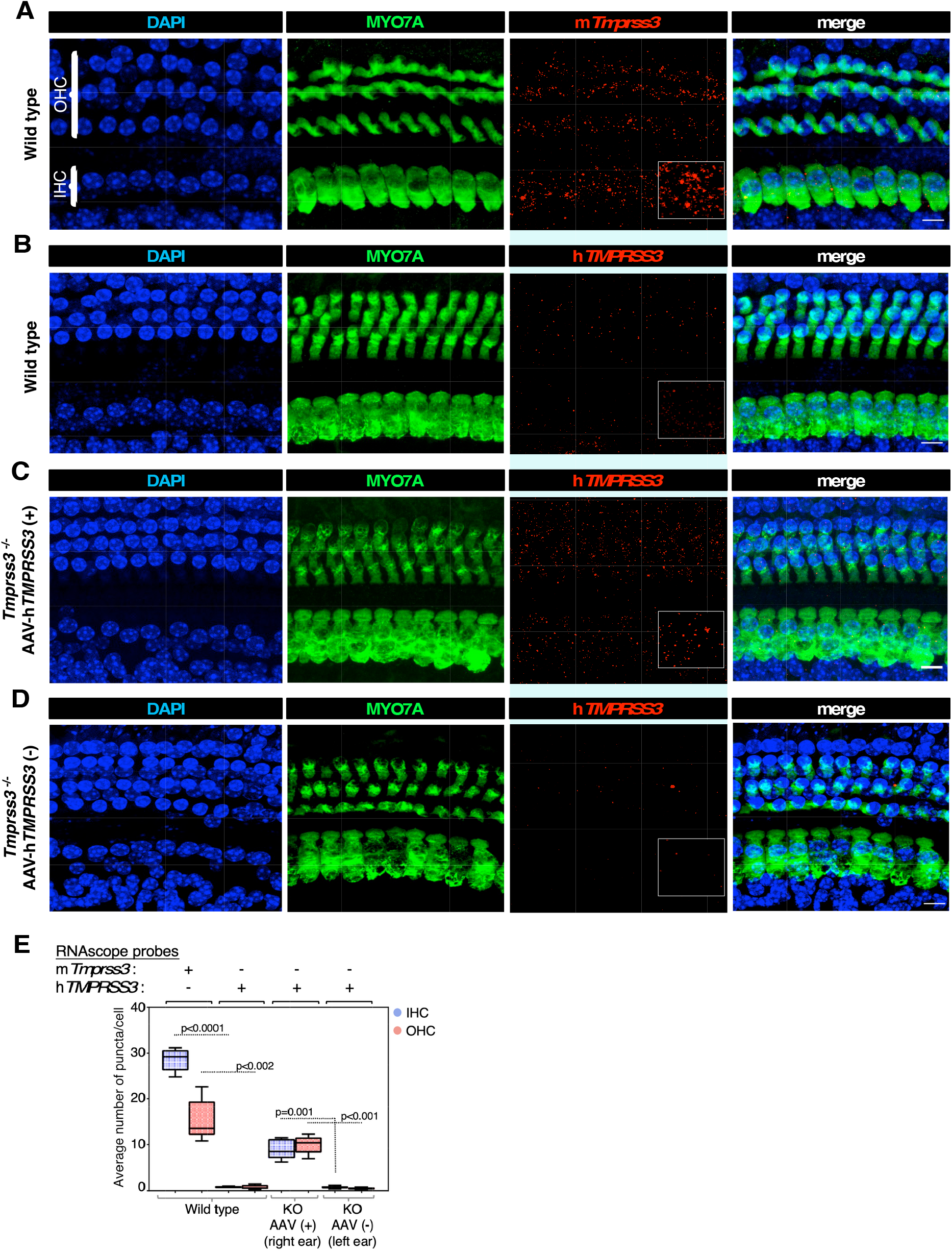
AAV2-*hTMPRSS3* inner ear delivery recapitulated endogenous *Tmprss3* expression in hair cells in adult *Tmprss3*^*A306T/A306T*^ KI mice. (A) Endogenous mouse *Tmprss3* gene expression in adult WT cochlea was detected by a mouse *Tmprss3* probe using RNAscope in a pattern overlapping with the inner and outer hair cells (IHC and OHC). (B) In adult WT cochlea, a human *TMPRSS3* probe showed very low or no cross-reactivity with the endogenous mouse *Tmprss3* mRNA. (C) In adult *Tmprss3*^*A306T/A306T*^ cochlea injected with AAV2-*hTMPRSS3*, a human *TMPRSS3* probe detected the transgene expression of the human *TMPRSS3* mRNA in the IHC and OHC. (D) In adult *Tmprss3*^*A306T/A306T*^ cochlea without injection, a human *TMPRSS3* probe did not detect mRNA above the background by RNAscope. Positive signals by RNAscope are shown as red punctuates in black background. Hair cells were labeled with MYO7A. The white square marked the area with a higher magnification for better visualization. Scale bars: 10 μm. (E) Quantification of mouse Tmprss3 or human *TMPRSS3* mRNA per inner and outer hair cell by RNAscope. Data are from 5 z-stack images obtained from the apex and/or apex-mid turn regions of each sample group (control n=3, AAV n=2). Results are expressed as an average number of dots per inner and outer hair cell and are displayed as a scatter column plot with medians indicated by the horizontal bar. P values are shown on top of each dashed line between compared groups.

The failure of cochlear implant in some *TMPRSS3* patients suggests the spiral ganglion neurons were affected by *TMPRSS3* mutations. However, it is not known if *TMPRSS3* is expressed in the adult ganglion neurons and any defect in the ganglion neurons is the direct result of the lack of *TMPRSS3* expression in the neurons or an indirect result of other cells that lack *TMPRSS3* expression. We examined the RNAscope data in the modiolus region that houses the spiral ganglion neurons. RNAscope showed a distinct expression pattern of endogenous *Tmprss3* mRNA by the mouse *Tmprss3* probe in the modiolus region with the level lower than in hair cells (**Figure 4A**). Combining with the immunofluorescence staining of HuD proteins to visualize spiral ganglions’ soma, the *Tmprss3* expression largely overlapped with the spiral ganglion neurons (**Figure 4A**). Signals were also detected in the region outside the spiral ganglions, suggesting that *Tmprss3* is expressed in other cell types in the modiolus region as Schwann cells and satellite cells (**Figure 4A**). Again, the human *TMPRSS3* probe did not show the signals above the background in the WT mouse modiolus region (**Figure 4B**), consistent with the specificity of the probe against human *TMPRSS3 mRNA*.

**Figure. 4.**
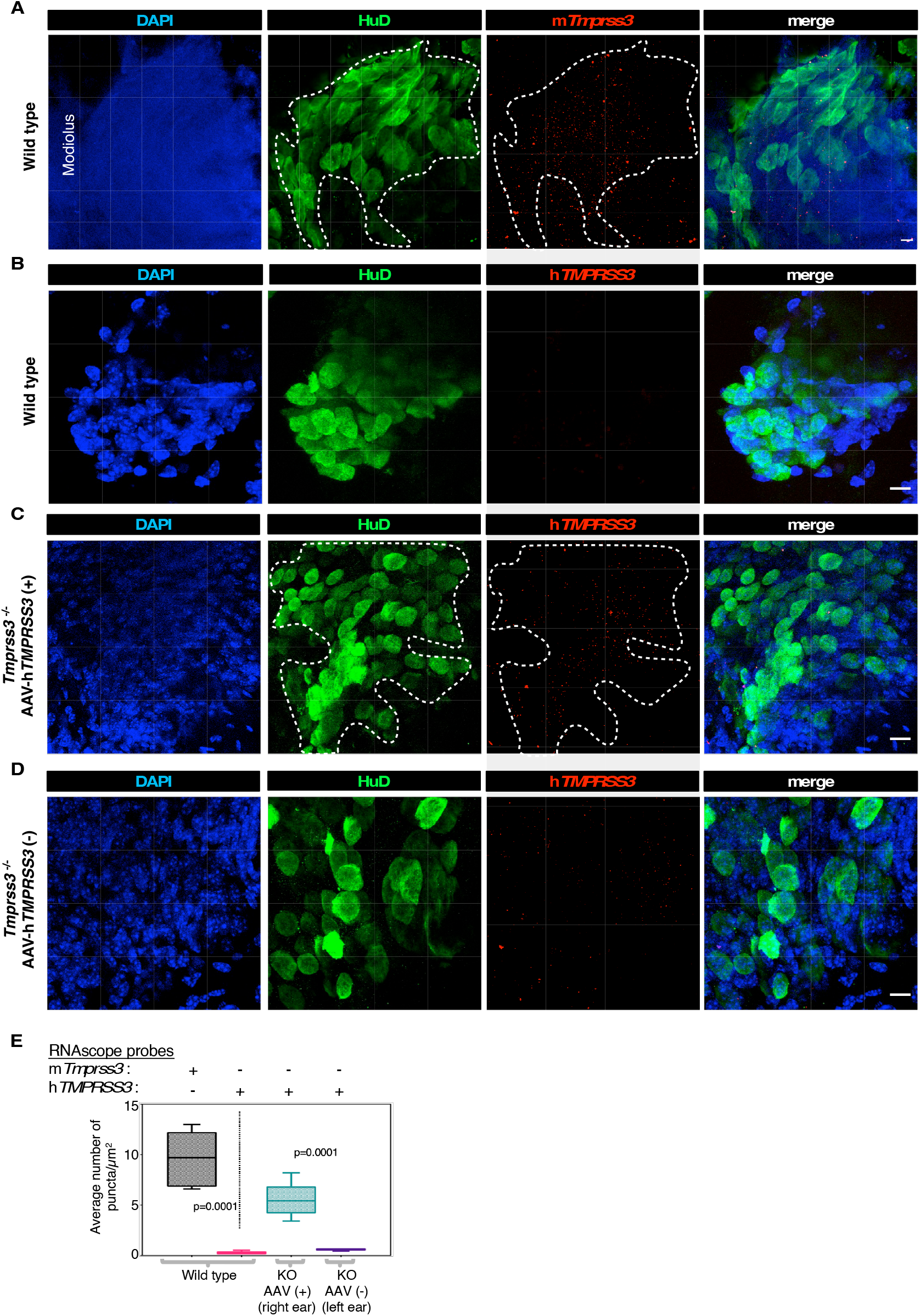
AAV2-*hTMPRSS3* inner ear delivery recapitulated endogenous *Tmprss3* expression in spiral ganglion neurons in adult *Tmprss3*^*A306T/A306T*^ KI mice. (A) In the adult WT mouse modiolus, endogenous mouse *Tmprss3* gene expression was detected by a mouse *Tmprss3* probe using RNAscope in a pattern largely overlapping with the spiral ganglion neurons (HuD, dotted cycle). (B) In adult WT modiolus, a human *TMPRSS3* probe showed no cross-reactivity with the endogenous mouse *Tmprss3* mRNA. (C) In adult *Tmprss3*^*A306T/A306T*^ modiolus injected with AAV2-*hTMPRSS3*, a human *TMPRSS3* probe detected the transgene expression of the human *TMPRSS3* mRNA in the spiral ganglion neurons (HuD, dotted cycle). (D) In adult *Tmprss3*^*A306T/A306T*^ modiolus without injection, a human *TMPRSS3* probe did not detect mRNA above the background by RNAscope. Scale bars: 10 μm. (E) Quantification of mouse *Tmprss3* or human *TMPRSS3* mRNA in modiolus region from the above-described samples. Data are from 5 z-stack images obtained from the apex and/or apex-mid turn regions of each sample group (control n=3, AAV n=2). Results are expressed as an average number of dots per modiolus and are displayed as a scatter column plot with medians indicated by the horizontal bar. P values are shown on top of each dashed line between compared groups.

We analyzed the RNAscope results of the human *TMPRSS3* probe hybridized to the *Tmprss3*^*A306T/A306T*^ inner ear injected with AAV2-*hTMPRSS3*. We detected the expression of the human *TMPRSS3* transgene in the modiolus that overlapped with the spiral ganglion neurons in AAV2-*hTMPRSS3* injected *Tmprss3*^*A306T/A306T*^ inner ear (**Figure 4C**), which coincided with the *Tmprss3* expression pattern in WT mice, supporting that AAV2 targeted the spiral ganglion neurons. In uninjected contralateral *Tmprss3*^*A306T/A306T*^ inner ear, no *TMPRSS3* expression was detected above the background (**Figure 4D**). Quantification confirmed that the human *TMPRSS3* gene was expressed, although at a level lower than the endogenous *Tmprss3* expression (**Figure 4E**). The RNAscope study showed that the endogenous *Tmprss3* is expressed in the hair cells and spiral ganglion neurons in adult WT inner ear, and AAV2-*hTMPRSS3* mediated gene delivery targeted the same cell population in the *Tmprss3*^*A306T/A306T*^ inner ears with the expression of the human *TMPRSS3* gene.

### AAV2-h*TMPRSS3 g*ene therapy rescues hearing in *Tmprss3*^*A306T/A306T*^ mice

As hearing loss in *Tmprss3*^*A306T/A306T*^ mice becomes apparent at 16.5 months of age, we performed inner ear AAV2-h*TMPRSS3* injection by the route of the round window with canal fenestration^52^ in *Tmprss3*^*A306T/A306T*^ mice aged 12 to 21 months, tested hearing one month after injection and continued the testing monthly for five months. Uninjected contralateral control ears served as controls. WT CBA/CaJ mice at the comparable ages were tested as additional controls to show the normal hearing profile in the mouse strain over time. Representative ABR wave forms recorded from a *Tmprss3*^*A306T/A306T*^ homozygous mutant mouse ear, an AAV2-*hTMPRSS3* treated *Tmprss3*^*A306T/A306T*^ ear and a WT mouse ear were shown 2 months after injection at the age of 20.5 months, using 11.32kHz tone bursts at incrementally increasing sound-pressure levels from 20dB to 90dB (**Figure S3A**). One month after injection, overall lower ABR thresholds were detected in injected ears compared to uninjected control ears at the frequencies of 5.66, 16 and 32 kHz. For the frequencies below 45.42 kHz, the ABR threshold reduction ranged from 6 dB at 11.32 kHz to 16 dB at 16 kHz, with an average reduction of 11 dB. No difference of DPOAE thresholds were detected between injected and uninjected ears (**Figure S3B**). The wave 1 amplitudes were statistically larger at 32 kHz above 50 dB SPL (**Figure S3G**). Two months after injection, ABR thresholds were further reduced at all frequencies in injected ears compared to uninjected control ears, ranging from 8 dB at 8 kHz to 18 dB at 45.24 kHz, with an average reduction of 15 dB across all frequencies (**Figure 5B**). Generally lower DPOAE thresholds were detected in injected compared to uninjected ears, with a significant reduction of 18 dB at 45.24 kHz (**Figure S3C**). At 3 and 4 months after injection, significant reductions in ABR thresholds were persistent in all frequencies in injected ears (**Figure 5C, D**), with an average reduction of 15 dB and 14 dB at 3 and 4 months, respectively. At the two time points, the DPOAE thresholds were generally lower and the wave 1 amplitudes at 32 kHz were larger in injected ears than uninjected control ears (**Figure S3D, E and 3I, J**). By 5 months post injection, with the exception of 16 kHz, ABR thresholds were still lower by an average of 9 dB (**Figure 5E**). Significant reduction in the DPOAE thresholds was detected at 8, 11.32 and 45.24 kHz (**Figure S3F**) and larger but not significant wave 1 amplitudes (**Figure S3K**) were seen in injected ears. At this age, the ABR thresholds in injected ears were indistinguishable from that of WT background CBA/CaJ strain at a comparable age of 23.5 months, which was elevated from that of CBA/CaJ of 22.5 months of age, an indication of age-related hearing loss in the CBA/CaJ background strain. In hearing tests of all ages, there was no significant difference of ABR thresholds between injected and WT CBA/CaJ ears (**Figure 5A-E**), a demonstration of robust rescue of hearing by AAV2-*hTMPRSS3* delivery in the *Tmprss3*^*A306T/A306T*^ ears to a level similar to WT control ears. Based on the data we conclude that a single administration of gene therapy by AAV2-*hTMPRSS3* results in long-term hearing rescue in the *Tmprss3*^*A306T/A306T*^ ears.

**Figure. 5.**
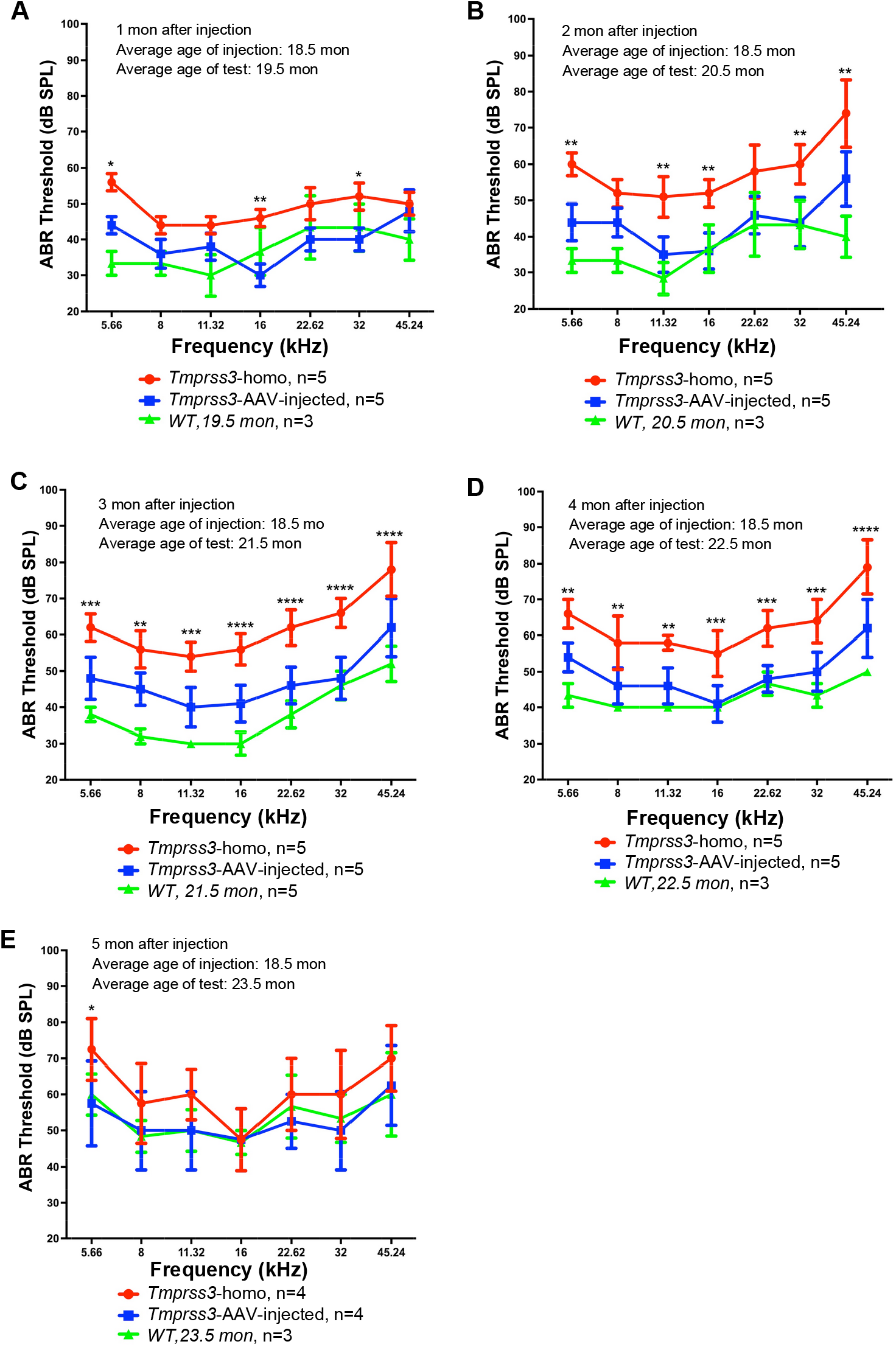
AAV2-h*TMPRSS3* injection in older *Tmprss3*^*A306T/A306T*^ mice rescues auditory function. (A-E) ABR thresholds in *Tmprss3*^*A306T/A306T*^ homozygous mice treated with AAV2-h*TMPRSS3* ears (blue), untreated *Tmprss3*^*A306T/A306T*^ homozygous ears (red), and wild-type (WT) CBA/CaJ mice (green) at one month after injection (A), two months after injection (B), three months after injection (C), four months after injection (D), and five months after injection (E), respectively. Significant hearing rescue was detected at all the time points with the exception at 5 months post injection when WT CBA/CaJ mice started to exhibit hearing loss. Values and error bars reflect mean ± SEM. Statistical analyses were performed by two-way ANOVA with Bonferroni correction for multiple comparisons: P value style, <0.05 (*), <0.01 (**), <0.001(***), <0.0001(****).

### AAV2-h*TMPRSS3* promotes HCs and SGNs survival in *Tmprss3*^*A306T/A306T*^ mice

Expression of *Tmprss3* in hair cells and the spiral ganglion neurons suggests the requirement of TMPRSS3 in hair cells and spiral ganglion neurons. To study the effect of the *TMPRSS3* c.916G>A mutation on the inner ear and determine how AAV2-h*TMPRSS3* gene delivery rescued the cellular phenotype, we performed immunolabeling of injected and uninjected cochleae and quantified IHCs, OHCs, and spiral ganglion neurons 2 months after injection at 20.5 months. In uninjected *Tmprss3*^*A306T/A306T*^ inner ears, a loss of OHCs in the regions of base and middle-base was detected (**Figure 6A1-A3**). In contrast in injected *Tmprss3*^*A306T/A306T*^ ears, more OHCs survived in same regions (**Figure 6B1-B3, 6C**). No significant difference was detected in IHC number between injected and uninjected control ears although the IHC average was higher in the injected ears (**Figure 6A1-B3, 6D**). In the base to mid-base turns of the modiolus region of uninjected *Tmprss3*^*A306T/A306T*^ ears, the TuJ1 labeling was condensed and localized to one side of the spiral ganglion neurons (**Figure 6E1, E2**). In contrast normal evenly distributed cytoplasmic labeling of TuJ1 was detected in the spiral ganglion neurons of injected ears (**Figure 6F1, F2**). Further significantly more ganglion neurons of more than threefold increase were detected in injected ears compared to uninjected ears (**Figure 6G**). Taken together, these results demonstrate that the *TMPRSS3* c.916G>A mutation caused the loss of the outer hair cells and spiral ganglion neurons from base to mid-base at 20.5 months and AAV2-h*TMPRSS3* gene therapy rescued hair cell and spiral ganglion neurons in the same regions.

**Figure. 6.**
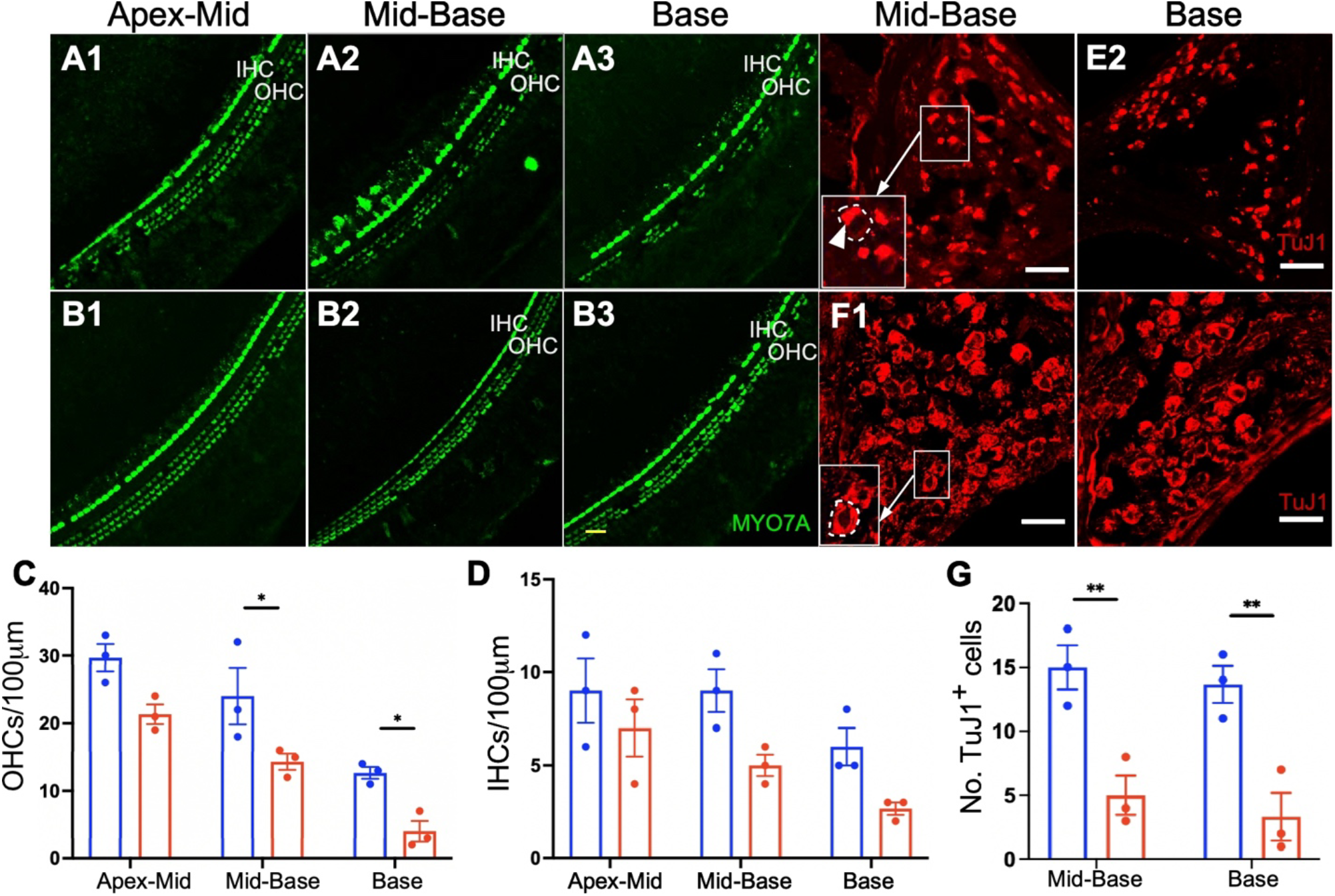
AAV2-h*TMPRSS3* injection rescues outer hair cells and spiral ganglion neurons in *Tmprss3*^*A306T/A306T*^ mice. (A-B) Representative confocal images of cochleae harvested at 20.5 months from uninjected (A) and AAV2-h*TMPRSS3* injected (B) ears of *Tmprss3*^*A306T/A306T*^ mice. The apex-middle (A1, B1), middle-base (A2, B2), and base (A3, B3) turns were dissected and stained with MYO7A (green) for hair cells. (C-D) Quantification and comparison of OHCs (C) and IHCs (D) in three representative regions of apex-middle, middle-base and base of injected (blue) and uninjected (red) *Tmprss3*^A306T/A306T^ cochleae from three mice at average of 20.5 months. Significantly more outer hair cells survived in injected ears compared to uninjected ears in the base and middle-base turns. (E-F) Representative confocal images of a modiolus cross section through the whole cochlea harvested at 20.5 months from uninjected (E) and AAV2-h*TMPRSS3* injected (F) ears of *Tmprss3*^*A306T/A306T*^ mice. The middle-base modiolus (E1, F1) and base modiolus (E2, F2) were stained with TuJ1 (red) for the spiral ganglion neurons. Arrows point to enlarged insets with higher magnifications of the square. Dotted circles showed an example of a spiral ganglion neuron soma with TuJ1 labeling, with condensed TuJ1 labeling (arrowhead) localized to one side within a SGN in uninjected ear (arrow). (G) Quantification and comparison of TuJ1 positive SGNs between uninjected (red) and AAV2-h*TMPRSS3* injected (blue) *Tmprss3*^*A306T/A306T*^ cochleae at average of 20.5 months. n=3. Values and error bars reflect mean ± SEM. Statistical analyses were performed by two-way ANOVA with Bonferroni correction for multiple comparisons: P value style, <0.05 (*), <0.01 (**), <0.001(***), <0.0001(****). Scale bars: 25μm.

## Discussion

With the goal to develop gene therapy for DFNB8 in humans, we created a knock-in mouse model with a frequent human DFNB8 *TMPRSS3* mutation that exhibits late onset and progressive hearing loss similar to DFNB5 patients. We show that one-time administration of AAV2-h*TMPRSS3* gene therapy in older age KI mice restores hearing at two years of age. Gene replacement therapy with the human *TMPRSS3* promotes the survival of hair cell and SGN, both are required for hearing and the latter is essential to treatment outcome of cochlear implants.

TMPRSS3 is a type II membrane serine protease and is associated with cancer development.^53,54^ *Tmprss3* has been found to be expressed in the developing cochlea including hair cells and ganglion neurons.^55^ The lack of TMPRSS3 has resulted in hair cell death in mouse organoids and ganglion neuron degeneration *in vitro*.^56,57^ Patients with homozygous *TMPRSS3* mutations manifest either postlingual progressive hearing loss (DFNB8) or congenital profound hearing loss (DFNB10) (https://hereditaryhearingloss.org/recessive-genes),^39,58^ likely due to the nature of the mutations. In *TMPRSS3* patients, cochlear implant is a standard form of treatment. Recently it is suggested in some *TMPRSS3* patients, the treatment outcome may diminish over time, leaving the patients without any treatment option.^67,71^ We aim to develop a gene therapy strategy to rescue hearing using a *TMPRSS3* mouse model with the potential to be further developed for the clinic.

AAV is one of the most effective gene delivery vectors that transduces both dividing and non-dividing cells to provide long-term gene expression.^59^ In the inner ear, prior studies found that conventional AAVs had good ability to transduce IHCs but there was little transduction in OHCs.^31,32^ We previously reported that AAV2 harboring GFP can transduce almost all IHCs and a large number of OHCs in adult C57BL/6 mouse cochleae by canalostomy.^60^ This delivery route was also used in CBA/CaJ mouse cochleae to show efficient IHC and OHC transduction.^61^ In a recent study, it was demonstrated that AAV2 transgene expression is in both OHCs (84%) and IHCs (97%) in adult C3H/FeJ mouse cochleae using a round window membrane plus canal fenestration delivery method.^62^ Combined, these studies have indicated that animal strain, animal age, viral titer, viral promoter, and even viral processing and purifying procedure can affect transduction efficiency.^52,63^ In this study, human *TMPRSS3* gene was delivered into a mouse model by AAV2 in CBA/CaJ background using a round window membrane with canal fenestration delivery route, which showed robust transduction of IHCs and efficient transduction of OHCs.

Gene therapy has been extensively used to treat mouse models for human genetic hearing loss with success.^13,19,25,26,33,64^ A majority of inner ear gene therapies however were only successful when performed in neonatal or young inner ears but not in adult.^13,23^ This could be due to the fact many types of genetic hearing loss are congenital and the inner ear cell types to be targeted are severely damaged or no longer available when mature, thus limiting the efficacy by intervention in mature inner ear. Further, the delivery vehicles may show difference targeting neonatal or adult cochlear cell types.^10,60,65^ In humans, even newborn inner ears are fully mature, any gene therapy strategy for patients would require the establishment of the window of opportunity for treatment and the demonstration that successful hearing rescue can be achieved in fully mature animal inner ears. *Tmprss3* knockout mouse exhibits severe congenital hair cell loss and profound hearing loss,^39^ thus is not a suitable model to develop *TMPRSS3* gene therapy to rescue hearing in mature inner ear. We chose to create a mouse model with a frequent founder human *TMPRSS3* mutation c.916G>A (p.A306T) that causes postlingual progressive hearing loss of DFNB8 in patients.^42,44^ To circumvent the mouse inbred strains such as C57 with accelerated age-related hearing loss, we chose to use CBA/CaJ as the background strain, so the strain-dependent hearing loss is not manifested until at an advanced age, which allowed us to study late onset progressive hearing loss and the treatment outcome in aged mice. The combination of mutation and mouse strain selections are important factors that enabled us to establish a knock-in mouse model of DFNB8 that is progressive HL. The model is ideal to test our gene therapy strategy to rescue hearing in not only mature but aged inner ear by one-time AAV administration.

It is surprising that the delivery of the mouse WT *Tmprss3* copy into normal adult mouse hair cells leads to severe hair cell loss and profound hearing loss. While the mechanism is not understood, it is likely that the level of the mouse TMPRSS3 protein is important to the survival of hair cells and essential for normal hearing. The detrimental effect of the mouse TMPRSS3 on hair cells and hearing prompted us to test the human *TMPRSS3* gene. Again, it is to our surprise that the human TMPRSS3, despite sharing a high degree of homology with the mouse TMPRSS3, 88% identical on the protein sequence (Figure S4)^66^, does not cause hair cell loss or hearing loss. Given our goal of using the human *TMPRSS3* gene for the clinical development, we have thus chosen the human *TMPRSS3* for gene therapy intervention in the KI TMPRSS3 mouse model.

Due to the lack of a suitable anti-TMPRSS3 antibody we carried out RNA-fluorescence *in situ* hybridization (RNAscope) to study endogenous *Tmprss3* expression in adult mouse inner ear and evaluate the inner ear cells targeted by AAV2-hTMPRSS3. In mouse *Tmprss3* has been shown to be expressed in hair cells, and other cochlear cell types including the cells in the modiolus region during development.^39,58^ A recent study also showed low *TMPRSS3* expression in human auditory neurons.^58^

By RNAscope, we found that AAV2 carrying the human *TMPRSS3* coding sequence can properly target IHCs, OHCs, and SGNs. The *TMPRSS3* transgene expression pattern resembles that of the WT *Tmprss3* expression. The data strongly support our approach of the AAV2 delivery in adult inner ear to target the cells that are directly affected by the lack of Tmprss3 expression and evaluate the rescue effect. A few studies have reported that the outcomes of cochlear implantation (CI) patients with *TMPRSS3*-related HL were inconsistent.^40,67-71^ Our results could provide a promising therapeutic avenue for those patients that have poor CI outcomes.

The mouse inner ear still undergoes development until the onset of hearing around P12.^72^ However, in the clinic, it is impractical to perform genetic therapies at the corresponding ages in human as the intervention would have to be carried out *in utero*. To translate these findings of preclinical mouse studies to humans, gene therapy research imperatively needs to focus on practical therapeutic strategies to perform in adult and aged mice. Among successful hearing rescue in animal models by gene or editing therapy, the vast majority of studies were carried out in the neonatal or infant stage.^13-17,19,21-25,29,32,35,73,74^ In *Tmprss3*^*A306T/A306T*^ mice, hearing loss does not start until ∼16.5 months of age, thus the intervention in aged mouse inner ears would be ideal to determine the treatment effect. We started injection in the animals of different ages, from 12 to 21 months, and measured hearing rescue temporally by the month post injection and with an average age at the injection time. Significant hearing rescue by the reduction in the ABR thresholds was detected one month after injection at some frequencies (**Figure 5A**). The rescue effect extended to all frequencies with increasing magnitudes (**Figure 5B-D**). Overall, there is a decrease in average DPOAE thresholds and greater wave 1 amplitudes in injected compared to uninjected contralateral control *Tmprss3*^*A306T/A306T*^ ears (**Figure S3**), supporting the improved outer hair cell function and neuronal activities. The improved hearing was detected 4 months after injection at an age of 22.5 months. At 5 months post injection with average age of mice at 23.5 months, even the WT control mice of the same age started to exhibit age-related hearing loss, with an ABR threshold average that was indistinguishable from injected *TMPRSS3*^*A306T/A306T*^mice (**Figure 5E**). We conclude that the strain specific age-related hearing loss (ARHL) obscured the treatment effect by AAV2-h*TMPRSS3* injection at 23.5 months. Our study is the first to demonstrate that one time administration of AAV gene therapy in aged mice is sufficient to rescue hearing long term in mice due to a *TMPRSS3* mutation. Hearing rescue by local injection at an advanced age in mice offers a wide therapeutic window that could enable the intervention in patients with DFNB8.

In the *Tmprss3*^*A306T/A306T*^ mice, significant hair cell especially OHC and neuronal loss was detected (**Figure 6**). In contrast to the *Tmprss3* knockout mice in which hair cells die rapidly^39^, our model with the A306T mutation showed a gradual hair cell and neuronal loss, which is consistent with late onset and progressive hearing loss. After AAV2-h*TMPRSS3* injection, both cell types survived significantly better, which is consistent with those cells targeted by AAV2. Interestingly, we also observed that TuJ1, a neuronal marker, showed an aggregation-like pattern with condensed TuJ1 signal restricted to neuron cell bodies unilaterally in the *Tmprss3*^*A306T/A306T*^ mice, which may further support previous findings showing cellular and molecular basis of SGN degeneration and the subsequent decline of the auditory nerve function in presbyacusis.^75^ While the physiological outcomes of TuJ1 aggregation is unknown, both the altered TuJ1 localization and the neuronal loss strongly support that direct impact of the *TMPRSS3* mutation A306T on the spiral ganglion neurons and the rescue effect by AAV2-h*TMPRSS3* local delivery. Mutations in *TMPRSS3* cause DFNB8 with hearing loss that is postlingual and progressive, and DFNB10 with congenital profound hearing loss. For DFNB8 patients, it is highly likely that hair cells and SGNs are present after birth, and they degenerate overtime. For DFNB8 patients, early intervention by AAV2-*TMPRSS3* gene therapy may be used as a standalone therapy to rescue hair cells and SGNs to prevent hearing loss. For *TMPRSS3* patients with profound hearing loss, their hair cells may have severely degenerated, and CI is the only option for treatment. For those patients, AAV2-*TMPRSS3* gene delivery can be used to promote the SGN survival to enhance CI treatment outcome long term. In addition to hearing loss caused by homozygous or compound heterozygous *TMPRSS3* mutations, there has been an increase in reports that heterozygous *TMPRSS3* mutations could contribute to accelerated ARHL^76^ in conjunction with a compound heterozygous mutation in another deafness gene.^77,78^ It is thus conceivable that gene therapy for *TMPRSS3* developed in this study could be expanded to rescue hearing in some aged patients with ARHL that represent the largest hearing loss population. The study supports that a single administration of AAV gene therapy for *TMPRSS3* in fully mature and aged inner ear could achieve noticeable restoration of hearing with long-term therapeutic effect. It is the proof of principle demonstration that gene therapy could be successfully implemented at a late stage in life and builds a solid foundation for its future clinical application.

## Methods Animals

*Tmprss3*^*A306T/A306T*^ mutant mice were generated at the Mouse Genome Engineering Core Facility of University of Nebraska Medical Center. The background was CBA/CaJ strain (Jaxson laboratories stock number 000656) to eliminate the effect of age-related hearing loss. The mice were housed in groups of two to five per cage and allowed free access to food and water. The animals were maintained under standard conditions (room temperature: 22 ±2°C; relative humidity: 55 ±10%) on a light: dark cycle of 12:12 h (6:00 a.m. to 6:00 p.m.). All procedures were approved by the Massachusetts Eye & Ear IACUC committee (protocol # 0804008) and University of Miami Institutional Animal Care Committee (protocol # 19-104), following the National Institutes of Health (NIH) Guidelines. All mouse experiments were performed in accordance with NIH guidelines for use and care of laboratory animals and were approved by the Massachusetts Eye & Ear IACUC committee (protocol # 0804008).

### AAV virus production

AAV2-m*Tmprss3* and AAV2-h*TMPRSS3* were produced by Kansas University Medical Center.

### Animal surgery

*Tmprss3*^*A306T/A306T*^ and wild type (WT) mice of either sex were anesthetized using intraperitoneal (i.p.) injection of ketamine (100mg/kg) and xylazine (10mg/kg). The post-auricular incision was exposed by shaving and disinfected using 10% povidone iodine. The AAV2-*Tmprss3* and AAV2-*hTMPRSS3* were injected into the inner ears of WT and *Tmprss3*^*A306T/A306T*^ mice. The AAV2-GFP was injected into the inner ears of WT mice. The total volume for each injection was 1 μl virus per cochlea.

### Auditory brainstem response and distortion product otoacoustic emission

Mice of either sex were anesthetized under the same conditions as for surgery. For ABR measurements, subcutaneous needle electrodes were inserted at the vertex (reference), ventral edge of the pinna (active electrode), and a ground reference near the tail. In a sound-proof chamber, mice were presented with 5-ms tone pips (delivered at 35/s). The response was amplified 10,000-fold, then filtered (100Hz-3kHz band-pass), digitized, and averaged (1,024 responses) at each SPL. The sound level was elevated in 5 dB steps from 20dB up to 90dB SPL at stimuli of 5.66-45.24 kHz frequencies (in half-octave steps). The “threshold” and wave 1 amplitude were identified as described previously.^17^ During the same recording session, DPOAEs were measured under the same conditions as for ABRs. Briefly, two primary tones (f2/f1=1.2) were set with f2 varied between 5.66 and 45.24 kHz in half-octave steps. Primaries were swept from 20dB SPL to 80dB SPL (for f2) in 5-dB steps. Thresholds required to produce a DPOAE at 5dB SPL were computed by interpolation as f2 level.

### Confocal microscopy and cell counting

Injected and uninjected cochleae were harvested from mice euthanized by CO2 inhalation. Temporal bones were fixed in 4% paraformaldehyde at 4°C overnight, then decalcified in 120mM EDTA for at least seven days until the tissues softened. After decalcification, the cochleae were dissected for whole mount immunostaining or cryosection at 10-μm thickness using published methods.^17,79,80^ Tissues were infiltrated with 0.25% Triton X-100 and blocked with donkey serum (5%) for one hour at room temperature, followed by washing 3 × 10 min with PBS and then incubated with primary antibody. Rabbit anti-MYO7A (1:500; #25-6790, Proteus BioSciences), mouse anti-Beta III tubulin (1:250; #801201, BioLegend), and chicken anti-GFP (1:1000; ab13970, Abcam) were used overnight. Tissues were then washed for 3 × 10 min with PBS and the secondary antibodies were incubated for one hour (Donkey anti-rabbit IgG Alexa Fluor Plus 488, 1:1000; Donkey anti-mouse IgG Alexa Fluor Plus 594, 1:1000; Thermo Fisher). Following secondary antibody incubation, tissues were washed for 3 × 10 min with PBS. Finally, tissues were placed on a microscope slide, and mounted with VECTASHIELD antifade mounting medium containing DAPI (VECTOR LABORATORIES, #H-1200). Images were taken with a Leica SP8 confocal laser scanning microscope (Leica Microsystems, Germany) via a 20X or 63X glycerin-immersion lens. Images were edited by ImageJ software and tools in ImageJ were used for counting of hair cells and SGNs. For hair cell counting, Myosin7a-positive hair cells per 100μm length were calculated in the apical, middle, and basal turns of cochleae. We counted at least two 100μm segments from at least three independent cochleae. For SGN cell counting, TUJ1-positive cells were calculated in the SGN area. The average cell number per 10^4^-μm^2^ was calculated for data analysis.

### RNA-FISH, immunohistochemistry, and data quantification

RNA-FISH was performed using RNAscope Multiplex Fluorescent Reagent Kit v2 (ACD, #323110), and the protocol described here is adapted from Huang and Eckrich with minor modifications.^50,51^ Briefly, membranous labyrinths were removed, and cochleae were immediately soaked in ice-cold 4% PFA in PBS. Cochleae were fixed for 2 hrs at RT on a shaker and washed for 3 × 10 min in 0.1% Tween-20 in PBS (PBT20), and subsequently dehydrated in a graded MeOH series (50, 75, and 100% in PBT20, 10 min for each grade). At the same time, probes were pre-warmed to 40°C for 10 min to allow aggregates to dissolve and then cooled down to RT. Protease III solution and Amplifiers I-IV were allowed to equilibrate to RT.

Before hybridization, cochleae were rehydrated at RT in a reverse MeOH series (100, 75, and 50% in PBT20; 10 min for each grade) and washed 6 × 5 min in PBT20. During the last washing step, the apical turn was dissected in PBT20 in a 35-mm sterile dish, and the stria vascularis and spiral ligament were carefully removed. Reissner’s membrane and the tectorial membrane were also separated from the sensory epithelium with fine forceps while the spiral ganglion was maintained intact. Dissected tissues collected from different sample groups were placed in individual mini cell strainers (pluriStrainer Mini 40 μm) containing 1 ml of PBT20, which were then transferred manually between the wells of a 24-well plate during all incubation and washing steps.

Dissected cochlea pieces were digested for 12 min at RT in 300 ml of Protease III solution. Following washing with 1 ml of PBT20 for 6 × 5 min to remove residual proteases, samples were incubated with hybridization probes for 2 hrs at 40°C. We used target probes against mouse *TMPRSS3* mRNA (ACD; #553861) or human *TMPRSS3* mRNA (ACD; #524691-C2). Subsequently, tissues were washed for 6 × 5 min with 1 ml of ACD washing buffer, re-fixed for 10 min with 4% PFA, and washed again for 6 × 5 min. Probe signals were amplified by incubation at 40°C in 4-5 drops of Amp1 (35 min), Amp2 (20 min), Amp3 (35 min), and Amp4 “Alt-A” solution (20 min), respectively. Following each step, tissues were washed for 6 × 5 min in 1 ml of ACD washing buffer in mini cell strainers housed in 24-well.

For a more precise identification of different cell types, the RNAscope assay was coupled to immunohistochemistry. Briefly, tissues were permeabilized for 10 min with 0.5% Triton X-100 in PBS (PBST), and blocked for 1 h with 10% donkey serum in PBST at RT. Subsequently, samples were incubated with primary antibodies against MYO7A (polyclonal rabbit, 1:250, #25-6790, Proteus Biosciences) and HuD (monoclonal mouse, 1:100, #sc-48421, SantaCruz) in antibody incubation buffer (5% donkey serum and 0.25% Triton X-100 in PBS) for 1 h at RT. Following washing with 0.1% Triton X-100 in PBS for 3 × 5 min, tissues were incubated at RT for 1h with appropriate secondary antibodies (Donkey α-rabbit IgG Alexa Fluor Plus 594, 1:500; Donkey α-mouse IgG Alexa Fluor Plus 647, 1:500; Thermo Fisher) in the antibody incubation buffer. Following secondary antibody incubation, tissues were washed for 3 × 5 min with 0.1% Triton X-100 in PBS. Finally, tissues were placed on a microscope slide, and mounted with Prolong Gold DAPI antifade medium (Invitrogen, #P36931), and allowed to penetrate and solidify overnight before imaging. RNAscope plus IHC experiments were repeated in more than two animals per group. For each sample group, ≥ five z-stack images were acquired using a 63X oil objective with 2X digital zoom in Leica SP8 confocal laser scanning microscope. Fluorescently labeled mRNA was quantified using the particle analysis tool in ImageJ, which requires 8-bit grayscale images. After adjusting thresholds to remove background signal for each image, the average number of mRNA molecules per hair cell was determined by dividing the number of particles by the number of MYO7A-positive hair cells or by the total number of particles per HuD-positive spiral ganglion neurons. For IHCs, OHCs, and SGNs counting, we acquired z-stacks by maximum intensity projections. The average number of IHCs and OHCs per hair cell was determined by dividing the number of particles by the number of MYO7A-positive hair cells or by the total number of particles per HuD-positive spiral ganglion neurons.

### Statistical Analysis

We used GraphPad Prism (v9, GraphPad Software, La Jolla, CA) for statistical analysis. For multiple comparisons in terms of ABRs and DPOAEs, statistical analyses were carried out by two-way ANOVA with Bonferroni corrections. For a numeric representation of RNAscope data, scatter column graphs were created using five z-stack images. For comparisons between two groups, data were analyzed by two-tailed unpaired t-tests with Welch’s correction. Significance threshold was set as p < 0.05 (p < 0.05 (*), p < 0.01 (**), p < 0.001(***), p < 0.0001(****)).

## ACKNOWLEDGEMENTS

This work was supported by U.S. NIH R01 DC016875, UG3TR002636, and U24HG010423 (to ZYC), R01 DC019404 (to XZL & ZYC), Ines-Fredrick Yeatts Fund (to ZYC.), R01DC012115 and R01DC005575 (to XZL).

## AUTHOR CONTRIBUTIONS

Conceptualization: WD, VE, HS, XZL, ZYC

Methodology: WD, VE, CL, MH, SS, AMA, ZH, CBG, HS, XZL, ZYC

Experiment: WD, VE, CL, MH, SS, AMA, ZH, HS

Data analysis: all authors Supervision: HS, XZL, ZYC

Writing—original draft: WD, VE, CL

Writing—review & editing: all authors

