## Supplemental Figures for "Rescue of Auditory Function by a Single Administration of AAV-*TMPRSS3* Gene Therapy in Aged Mice of Human Recessive Deafness DFNB8"

### Supplementary Figures

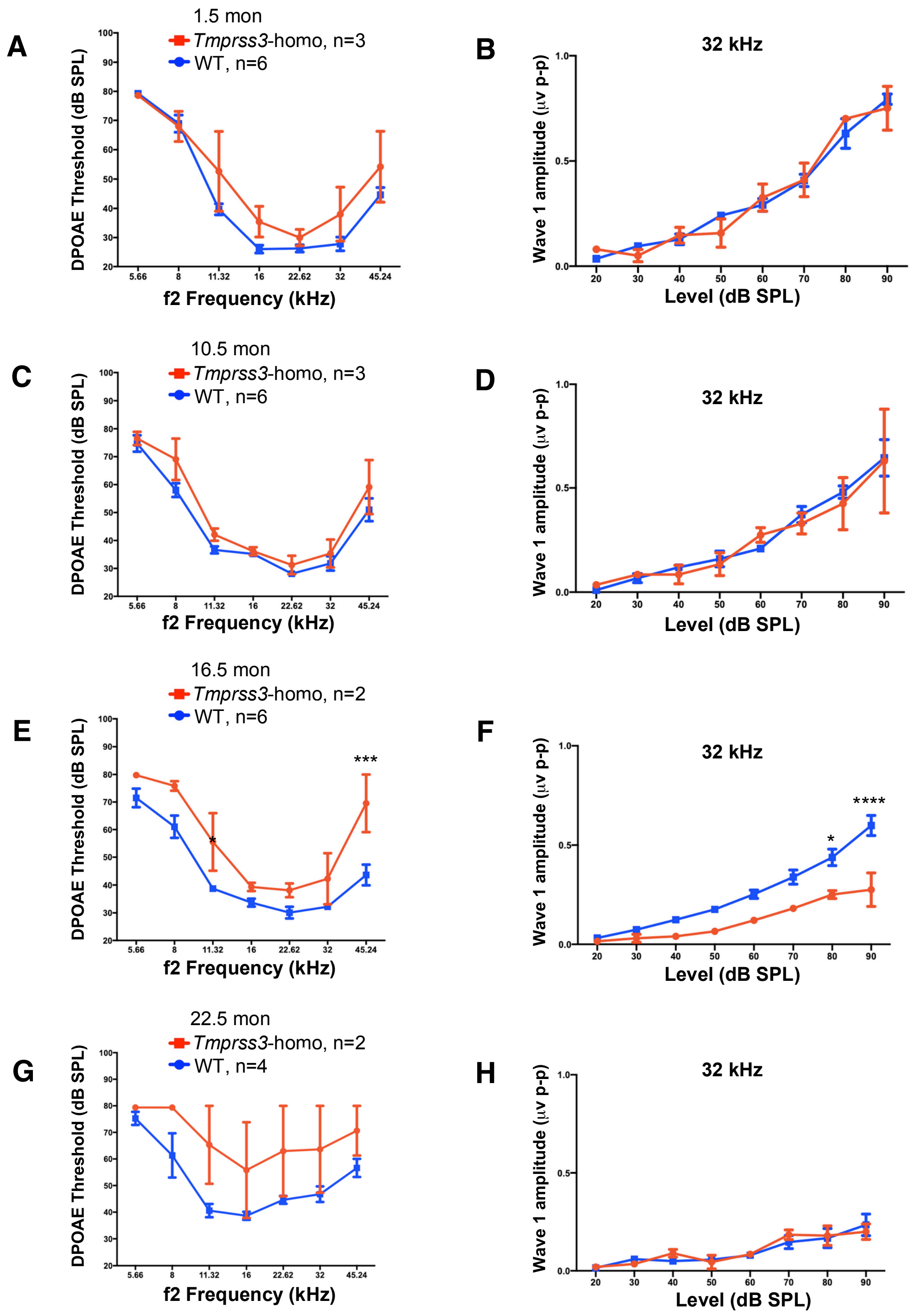

**Figure S1. *Tmprss3*$^{A306T/A306T}$ knock-in mouse model exhibits late-onset progressive hearing loss.** (A-D) Similar DPOAE thresholds and wave 1 amplitudes in *Tmprss3 Tmprss3*$^{A306T/A306T}$ homozygous mice ears (red) and wild-type ears (blue) at 1.5 months (A, B) and 10.5 months (C, D), respectively. (E) DPOAE thresholds were higher across frequencies in the *Tmprss3 Tmprss3*$^{A306T/A306T}$ homozygous ears (red) compared to WT ears at the similar age (blue), an indication of reduced outer hair cell function. (F) The wave 1 amplitudes were significantly higher at 32 kHz at 80 and 90 dB in WT ears compared to KI ears at 16.5 months. (G) At 22.5 months of age, DPOAE thresholds were higher across frequencies in the *Tmprss3 Tmprss3*$^{A306T/A306T}$ homozygous ears (red) compared to WT ears at the similar age (blue), although there was a big variation among the KI ears. (H) At 22.5 months, the wave1 amplitudes in WT and the KI ears were similar, which were lower than at 16.5 months, an indication that the age-related reduction in neural activities of WT ears that matched the reduction in the KI ears. Values and error bars reflect mean ± SEM. Statistical analyses were performed by two-way ANOVA with Bonferroni correction for multiple comparisons: P value style, <0.05 (*), <0.01 (**), <0.001(***), <0.0001(****).

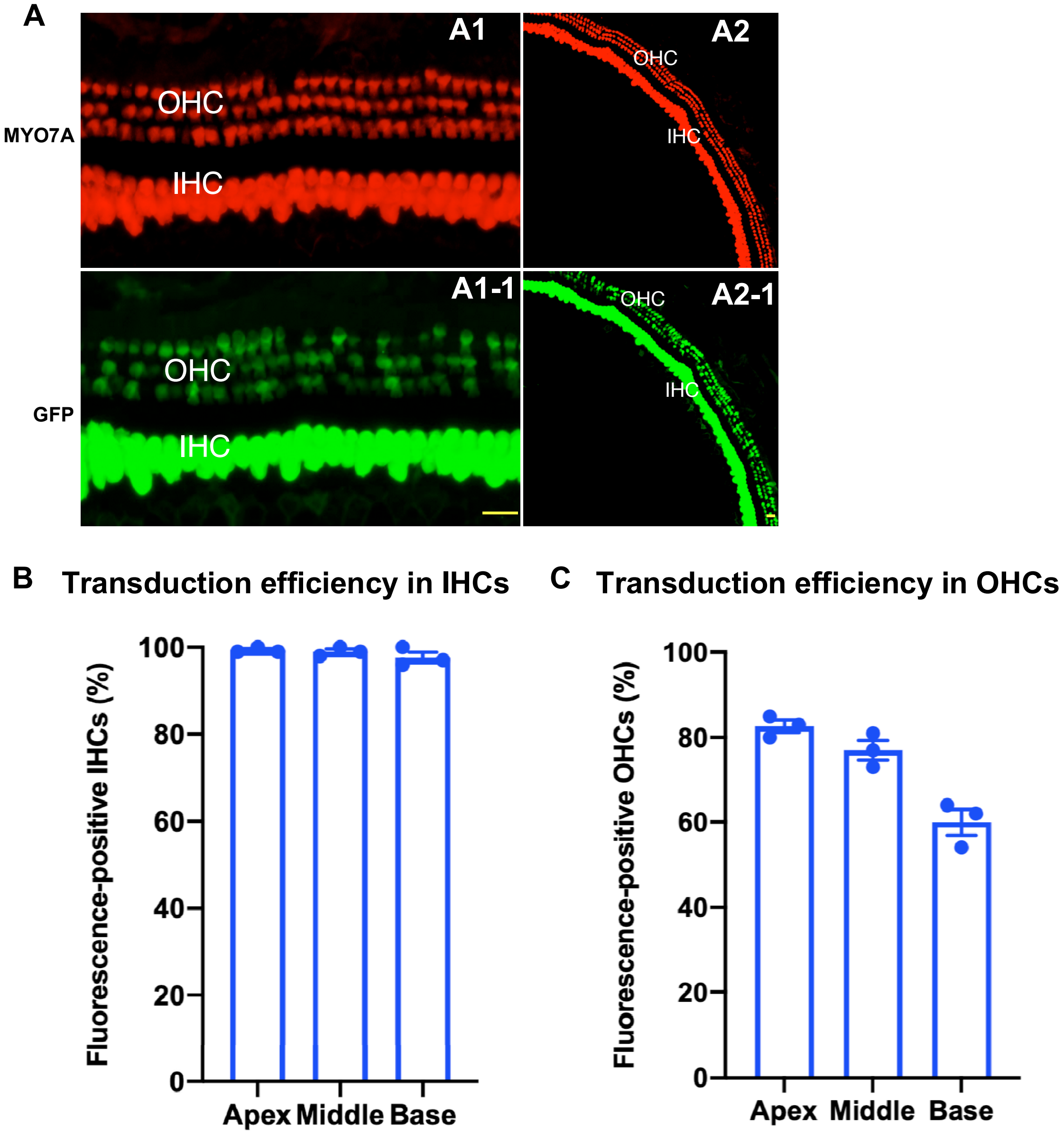
A
A1
A2
MYO7A
OHC
IHC
A1-1
A2-1
GFP
B
Transduction efficiency in IHCs
Fluorescence-positive IHCs (%)
100
80
60
40
20
0
Apex
Middle
Base
C
Transduction efficiency in OHCs
Fluorescence-positive OHCs (%)
100
80
60
40
20
0
Apex
Middle
Base

**Figure S2. AAV2 transduction in adult mice cochleae.** (A) Representative confocal images of apex-middle region of AAV2 transduction in adult mouse cochleae (a1, high magnification; a2, low magnification). Cochleae were stained with MYO7A (red, a1 and a2), anti-GFP (green, a1-1 and a2-1). Scale bar, 20μm. (B, C) Quantification of IHC (B) and OHC (C) AAV2-GFP transduction efficiency across different regions of the cochlea (apex, middle, and base) one month after injection at 5 months of age. Values and error bars reflect mean ± SEM, n=3.

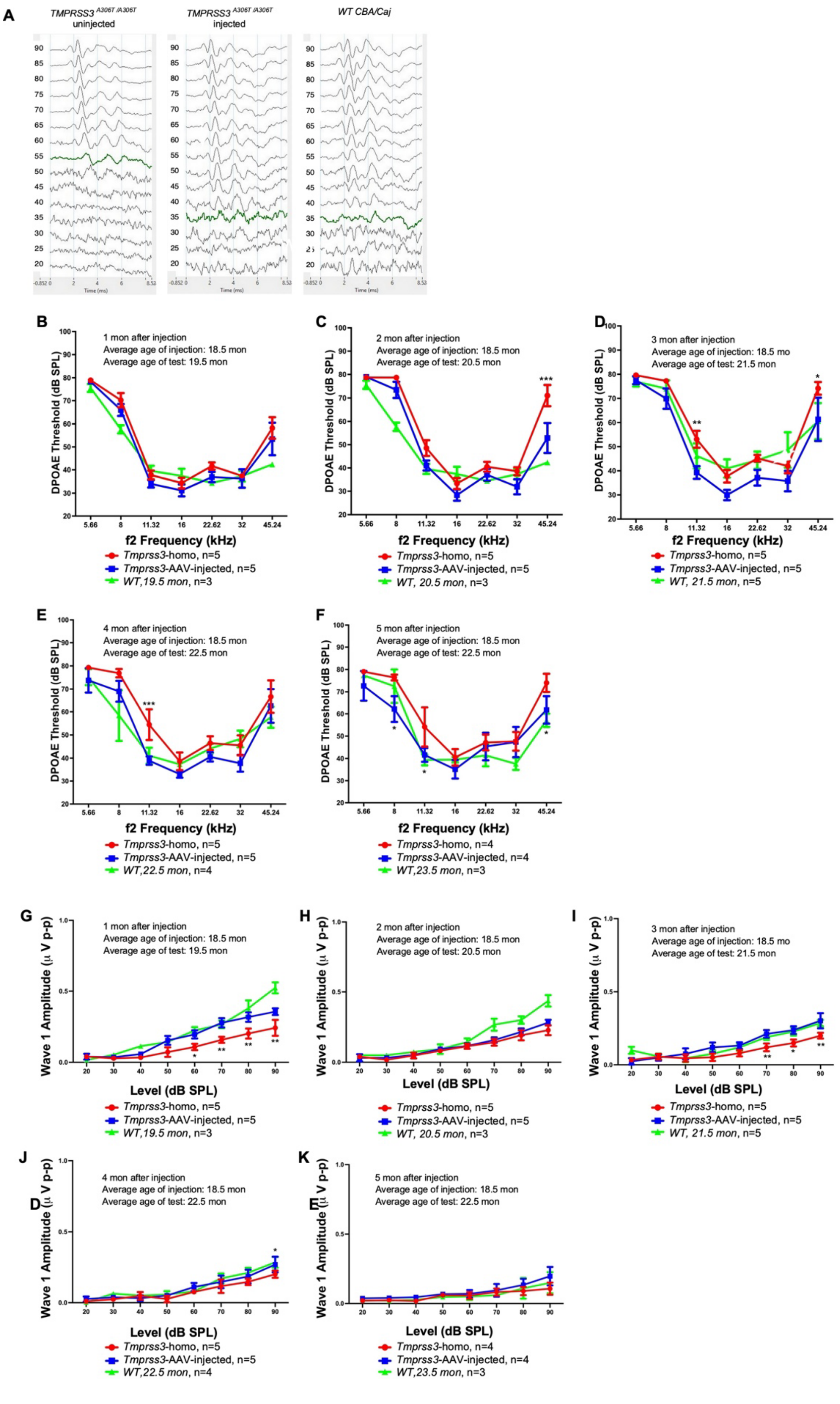
A
TMPRSS3 A306T/A306T uninjected
TMPRSS3 A306T/A306T injected
WT CBA/Caj
B
1 mon after injection
Average age of injection: 18.5 mon
Average age of test: 19.5 mon
DPOAE Threshold (dB SPL)
f2 Frequency (kHz)
Tmprss3-homo, n=5
Tmprss3-AAV-injected, n=5
WT,19.5 mon, n=3
C
2 mon after injection
Average age of injection: 18.5 mon
Average age of test: 20.5 mon
Tmprss3-homo, n=5
Tmprss3-AAV-injected, n=5
WT, 20.5 mon, n=3
D
3 mon after injection
Average age of injection: 18.5 mo
Average age of test: 21.5 mon
Tmprss3-homo, n=5
Tmprss3-AAV-injected, n=5
WT, 21.5 mon, n=5
E
4 mon after injection
Average age of injection: 18.5 mon
Average age of test: 22.5 mon
Tmprss3-homo, n=5
Tmprss3-AAV-injected, n=5
WT,22.5 mon, n=4
F
5 mon after injection
Average age of injection: 18.5 mon
Average age of test: 22.5 mon
Tmprss3-homo, n=4
Tmprss3-AAV-injected, n=4
WT,23.5 mon, n=3
G
1 mon after injection
Average age of injection: 18.5 mon
Average age of test: 19.5 mon
Wave 1 Amplitude (µ V p-p)
Level (dB SPL)
Tmprss3-homo, n=5
Tmprss3-AAV-injected, n=5
WT,19.5 mon, n=3
H
2 mon after injection
Average age of injection: 18.5 mon
Average age of test: 20.5 mon
Tmprss3-homo, n=5
Tmprss3-AAV-injected, n=5
WT, 20.5 mon, n=3
I
3 mon after injection
Average age of injection: 18.5 mo
Average age of test: 21.5 mon
Tmprss3-homo, n=5
Tmprss3-AAV-injected, n=5
WT, 21.5 mon, n=5
J
4 mon after injection
Average age of injection: 18.5 mon
Average age of test: 22.5 mon
Tmprss3-homo, n=5
Tmprss3-AAV-injected, n=5
WT,22.5 mon, n=4
K
5 mon after injection
Average age of injection: 18.5 mon
Average age of test: 22.5 mon
Tmprss3-homo, n=4
Tmprss3-AAV-injected, n=4
WT,23.5 mon, n=3

**Figure S3. AAV2-hTMPRSS3 transduction rescues auditory function in *Tmprss3$^{A306T/A306T}$* mice.** (A) Representative ABR waveform recorded from an untreated *Tmprss3$^{A306T/A306T}$* mouse ear (left), a *Tmprss3$^{A306T/A306T}$* ear treated with AAV2-*hTMPRSS3* ear (middle), and a WT ear (right) at 20.5 months using 11.32 kHz tone bursts, respectively. Threshold was determined by the detection of peak 1 and is indicated by green color trace. (B-F) DPOAE thresholds in *Tmprss3$^{A306T/A306T}$* mice treated with AAV2-*hTMPRSS3* ears (blue), untreated *Tmprss3$^{A306T/A306T}$* ears (red), and WT ears (green) at one month (B), two months (C), three months (D), four months (E), and five months after injection (F), respectively. Generally lower DPOAE thresholds were seen in injected compared to uninjected *Tmprss3$^{A306T/A306T}$* ears. (G-K) Amplitude of ABR wave 1 at 32kHz in *Tmprss3$^{A306T/A306T}$* mice treated with AAV2-*hTMPRSS3* ears (blue), untreated *Tmprss3$^{A306T/A306T}$* ears (red), and WT ears (green) at one month (G), two months (H), three months (I), four months (J), and five months after injection (K), respectively. Wave 1 amplitudes were generally higher in the injected compared to uninjected *Tmprss3$^{A306T/A306T}$* ears up to 4 months post injection. By 5 months post injection at 23.5 months, wave 1 amplitudes in WT, injected and uninjected KI ears were indistinguishable. Values and error bars reflect mean ± SEM. Statistical analyses were performed by two-way ANOVA with Bonferroni correction for multiple comparisons: P value style, <0.05 (*), <0.01 (**), <0.001(***), <0.0001(****).

```
UniProt|Q8K1T0|TMPS3_MOUSE      MGENDPPAAEAPFSFRSLFGLDDLKISPVAPDGDAVAAQILSLLPLKFFPIIVIGIIALI 60
UniProt|P57727|TMPS3_HUMAN      MGENDPPAVEAPFSFRSLFGLDDLKISPVAPDADAVAAQILSLLPLKFFPIIVIGIIALI 60
                                ********.***********************.***************************

UniProt|Q8K1T0|TMPS3_MOUSE      LALAIGLGIHFDCSGKYRCHSSFKCIELTARCDGVSDCKNAEDEYRCVRVSGQRAALQVF 120
UniProt|P57727|TMPS3_HUMAN      LALAIGLGIHFDCSGKYRCRSSFKCIELIARCDGVSDCKDGEDEYRCVRVGGQNAVLQVF 120
                                *******************:******** **********:.*********.**.*.****

UniProt|Q8K1T0|TMPS3_MOUSE      TAAAWRTMCSDDWKSHYAKIACAQLGFPSYVSSDHLRVDALEEQFQGDFVSINHLLSDDK 180
UniProt|P57727|TMPS3_HUMAN      TAASWKTMCSDDWKGHYANVACAQLGFPSYVSSDNLRVSSLEGQFREEFVSIDHLLPDDK 180
                                ***:*:********.***::**************:***.:** **: :****:*** ***

UniProt|Q8K1T0|TMPS3_MOUSE      VTALHHSVYMREGCTSGHVVTLKCSACGTRTGYSPRIVGGNMSSLTQWPWQVSLQFQGYH 240
UniProt|P57727|TMPS3_HUMAN      VTALHHSVYVREGCASGHVVTLQCTACGHRRGYSSRIVGGNMSLLSQWPWQASLQFQGYH 240
                                *********:****:*******:*:*** * *** ******** *:*****.********

UniProt|Q8K1T0|TMPS3_MOUSE      LCGGSIITPLWIVTAAHCVYDLYHPKSWTVQVGLVSLMDSPVPSHLVEKIIYHSKYKPKR 300
UniProt|P57727|TMPS3_HUMAN      LCGGSVITPLWIITAAHCVYDLYLPKSWTIQVGLVSLLDNPAPSHLVEKIVYHSKYKPKR 300
                                *****:******:********** *****:*******:*.*.********:*********

UniProt|Q8K1T0|TMPS3_MOUSE      LGNDIALMKLSEPLTFDETIQPICLPNSEENFPDGKLCWTSGWGATEDG-GDASPVLNHA 359
UniProt|P57727|TMPS3_HUMAN      LGNDIALMKLAGPLTFNEMIQPVCLPNSEENFPDGKVCWTSGWGATEDGAGDASPVLNHA 360
                                **********: ****:* ***:*************:************ **********

UniProt|Q8K1T0|TMPS3_MOUSE      AVPLISNKICNHRDVYGGIISPSMLCAGYLKGGVDSCQGDSGGPLVCQERRLWKLVGATS 419
UniProt|P57727|TMPS3_HUMAN      AVPLISNKICNHRDVYGGIISPSMLCAGYLTGGVDSCQGDSGGPLVCQERRLWKLVGATS 420
                                ******************************.*****************************

UniProt|Q8K1T0|TMPS3_MOUSE      FGIGCAEVNKPGVYTRITSFLDWIHEQLERDLKT 453
UniProt|P57727|TMPS3_HUMAN      FGIGCAEVNKPGVYTRVTSFLDWIHEQMERDLKT 454
                                ****************:**********:******
```

**Figure S4. Alignment of TMPRSS3 proteins from human and mouse.** Clustal Omega (v 1.2.4) alignment of TMPRSS3 proteins from human and mouse listed by their names and UniProt IDs showed that TMPRSS3 sequences are 88% identical between humans and mice. An asterisk (*) denotes identical residues; double dots (:) represent a conserved residue substitution; a single dot (.) shows partial conservation of the residue. Red, small, and hydrophobic; blue, acidic; magenta, basic; green, hydroxyl or sulfhydryl or amine or G.
